# Phylogeographic reconstruction using air transportation data and its application to the 2009 H1N1 influenza A pandemic

**DOI:** 10.1101/666982

**Authors:** Susanne Reimering, Sebastian Muñoz, Alice C. McHardy

**Author notes:** Corresponding author (ACM).

## Abstract

Influenza A viruses cause seasonal epidemics and occasional pandemics in the human population. While the worldwide circulation of seasonal influenza is at least partly understood, the exact migration patterns between countries, states or cities are not well studied. Here, we use the Sankoff algorithm for parsimonious phylogeographic reconstruction together with effective distances based on a worldwide air transportation network. By first simulating geographic spread and then phylogenetic trees and genetic sequences, we confirmed that reconstructions with effective distances inferred phylogeographic spread more accurately than reconstructions with geographic distances and Bayesian reconstructions with BEAST, the current state-of-the-art. Our method extends the state-of-the-art by using fine-grained locations like airports and inferring intermediate locations not observed among sampled isolates. When applied to sequence data of the pandemic H1N1 influenza A virus in 2009, our approach correctly inferred the origin and proposed airports mainly involved in the spread of the virus. In case of a novel outbreak, this approach allows to rapidly analyze sequence data and infer origin and spread routes to improve disease surveillance and control.

**Author summary:** Influenza A viruses infect up to 5 million people in recurring epidemics every year. Further, viruses of zoonotic origin constantly pose a pandemic risk. Understanding the geographical spread of these viruses, including the origin and the main spread routes between cities, states or countries, could help to monitor or contain novel outbreaks. Based on genetic sequences and sampling locations, the geographic spread can be reconstructed along a phylogenetic tree. Our approach uses a parsimonious reconstruction with air transportation data and was verified using a simulation of the 2009 H1N1 influenza A pandemic. Applied to real sequence data of the outbreak, our analysis gave detailed insights into spread patterns of influenza A viruses, highlighting the origin as well as airports mainly involved in the spread.

## Introduction

Influenza A viruses continue to impose high mortality and morbidity worldwide [1]. In addition to seasonal epidemics every winter, pandemics can occur when an antigenically novel virus, usually of zoonotic origin, establishes human-to-human transmission. The latest pandemic occurred in 2009, when a novel H1N1 influenza A virus emerged in March and quickly spread around the globe, with 177,000 confirmed infections in over 170 countries until early August [2]. While it is known that viral spread is mainly influenced by air travel [3] and seasonal epidemics are seeded from East and Southeast Asia [4], the exact migration patterns are not fully understood. Especially the inference of transition patterns on a fine-grained scale, e.g. between single countries, states or cities, remains a challenge.

If sequence data together with sampling locations are available, the origin and spread of viruses can be reconstructed using phylogeography. Given a phylogeny as well as locations for the leaf nodes of the tree, phylogeography infers locations for internal nodes of the tree. This approach reconstructs the source of the outbreak as well as spread routes. Current state-of-the-art methods for phylogeography are based on Bayesian inference. Discrete Bayesian phylogeography [5] followed efforts to use parsimonious methods for the reconstruction by minimizing the number of changes between states [6]. Bayesian methods improved this approach by incorporating uncertainty and branch lengths, giving posterior probabilities to evaluate the quality of the reconstruction and allowing the extension to a general linear model to test potential predictors of viral spread [3]. Therefore, Bayesian phylogeography is now commonly used to study viral pathogens like influenza [4], HIV [7] and Ebola [8]. However, these methods have several drawbacks that have not yet been addressed. First, due to the large number of parameters that are estimated in Bayesian phylogeographic studies, the analysis is slow for larger datasets and the number of distinct states is limited. With too many states compared to the number of sequences, the analysis cannot converge and estimate the rates due to a lack of data [9]. Instead, locations are often aggregated into larger regions such as continents [3,4], although more fine-grained locations such as countries, states or sometimes even cities are available for a lot of sequences. Second, these methods can only infer locations which are observed in the data, excluding possible intermediate states which have not been sampled. Continuous phylogeography, based on inferring geographic coordinates using Brownian diffusion models, are an alternative which allow to infer intermediate locations [10]. While this is a good model for local diffusion of rapidly evolving viruses, it is less applicable for viruses that travel both locally and over large distances in a very short time, e.g. by air travel in the case of influenza A viruses. Here, we propose a new parsimony-based approach for phylogeographic reconstruction and apply it to study the 2009 outbreak of the pandemic H1N1 (pH1N1) influenza A virus. To use prior knowledge about the mode of travel, we directly include air transportation data via effective distances, which are defined by the number of people travelling from one location to another [11]. The phylogeographic reconstruction then uses the Sankoff algorithm [12] to find internal locations, minimizing the distances along the tree. This approach inherently overcomes the shortcomings of the current state-of-the-art and allows both the use of fine-grained locations and the inference of intermediate locations.

We evaluated this approach using simulated data and our recently described distance measure to compare phylogeographic spread among different tree topologies [13]. We showed that effective distances calculated on air passenger flows yield more accurate reconstructions than geographic distances and Bayesian reconstructions with BEAST [14]. We then used this method to study the early spread of the pH1N1 influenza A virus. Our method correctly inferred Mexico as an origin. Further, we proposed a list of airports that were mainly involved in the initial spread of the virus and seeded a large number of infections in new locations. In the case of future pandemics, this method allows to quickly analyze viral sequence data to identify the origin and major spread routes, which could help to implement surveillance and control measures to contain the spread of the disease.

## Results

### Phylogeographic reconstruction using simulated data

The early spread of the pH1N1 influenza A virus was simulated using GLEAMviz [15], which has been widely used to simulate this outbreak [16,17] and has been shown to accurately predict influenza activity in various countries [18]. Based on the simulated transitions between locations during the first weeks of the pandemic, we then used FAVITES [19] to simulate the isolate sampling and sequencing, the tree as well as the sequences evolving along the phylogeny 50 times in total. The resulting simulated datasets included on average 97 sequences sampled from 76 unique locations in 23 countries. To confirm that the simulated sequences and the corresponding tree were an accurate representation of the pH1N1 virus, we compared them to real HA sequences of the outbreak that were sampled until the end of April 2009. We calculated pairwise distances between the sequences using a Jukes-Cantor model for both real and simulated sequences (S1 Fig A) and further inferred phylogenetic trees to compare branch length distributions (S1 Fig B). With both the genetic distances between sequences as well as branch lengths showing a similar distribution, we conclude that the simulation was an accurate representation of the pH1N1 influenza A virus in terms of sequence diversity and tree resolution, which is essential to achieve a comparable accuracy for phylogeographic reconstructions. Most sampling locations in the simulated data were in Mexico and the US, but the virus already spread to Canada, Europe, Asia and Oceania as well (Fig 1A). New infections were mainly seeded from Mexico, especially from Veracruz, the origin of the outbreak, and Cancún, the second largest airport of the country. Transitions were given via origin and destination, as output by the GLEAMviz simulation, but were not necessarily the direct route of travel. While Veracruz only has direct flights to a small number of locations, transitions to the US and Europe were possible via connecting flights, e.g. via Mexico City, the main connection from Veracruz. Overall, the simulated locations agreed with the spread of the pandemic until the end of April 2009, where the majority of cases was reported for Mexico and the US, and the first cases occurred on other continents as well [20].

**Fig 1.**
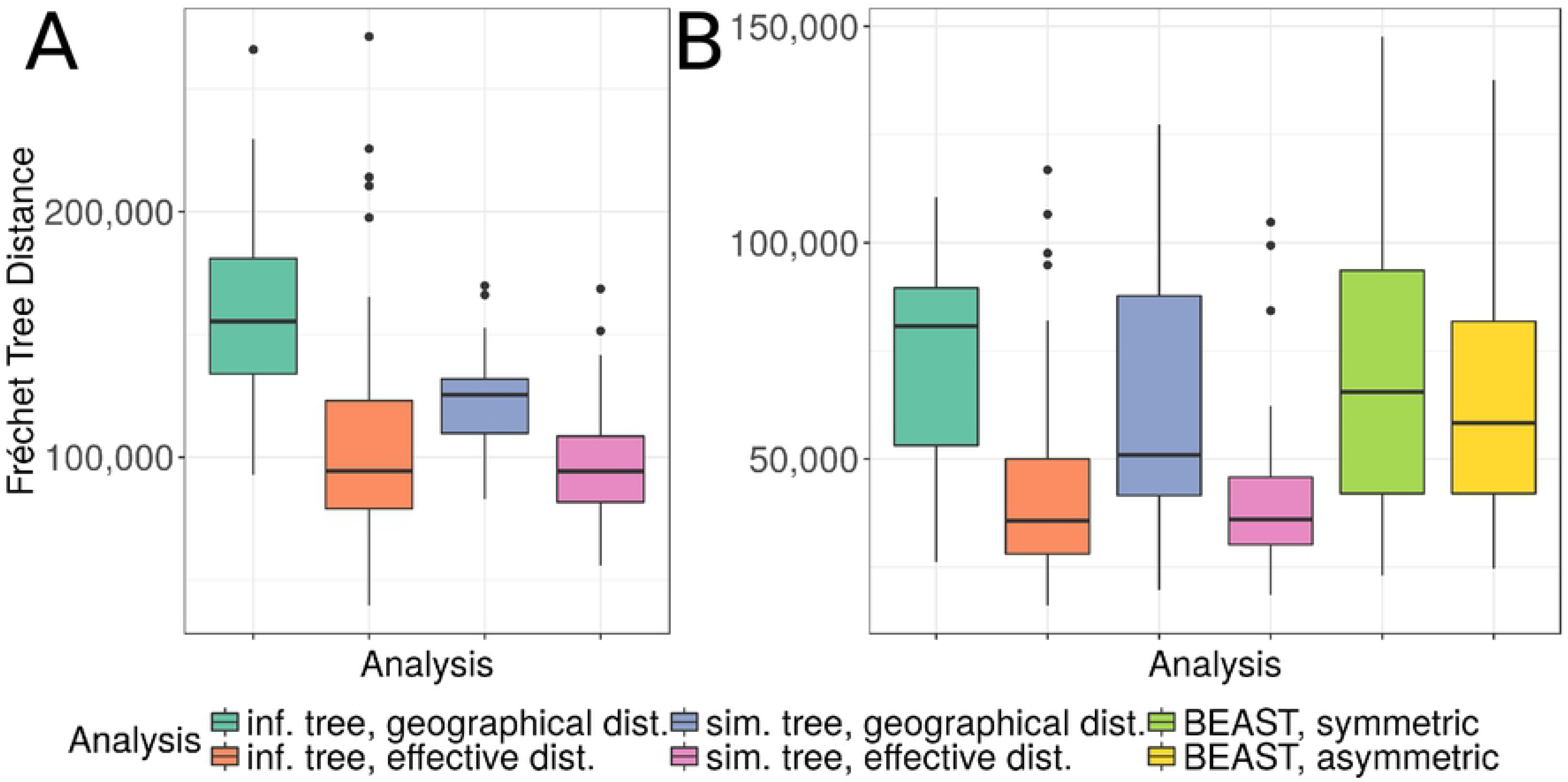
Simulated and reconstructed phylogeographic spread. Viral spread in North and Central America, shown by transitions between locations for the underlying ground truth of the simulation (panel A) and phylogeny, and the inferred spread and phylogeny in panel B, which was reconstructed based on sequences simulated along the phylogeny in A. For the phylogeographic reconstruction, effective distances were used. Transitions between locations are colored by their origin. In the simulation, the outbreak was set in Veracruz (yellow), which was mainly involved in the spread, together with Cancún (green). Transitions from Mexico City are shown in orange, all others in dark blue. In the reconstruction, the origin was placed in Zacatecas (light blue). From there, the virus mainly spread via Mexico City (orange), Cancún (green) and Chicago (pink). All other transitions are shown in dark blue. The actual origin in Veracruz was not inferred and no sequences were sampled here, which is why this location is missing in the map. While the origin and main spread routes differ, the locations are geographically close, leading to a Fréchet tree distance of 39,591.53. The spread was visualized using Spread3 [21].

For each simulated dataset, we used the tree inferred on the simulated sequences as well as their sampling locations for a phylogeographic reconstruction with the Sankoff algorithm (Fig 2). We tested both geographic and effective distances. Further, the phylogeographic reconstruction was done using the simulated tree topology to assess the level of variation introduced by the tree inference and the robustness of our method in case of inaccurate topologies.

**Fig 2.**
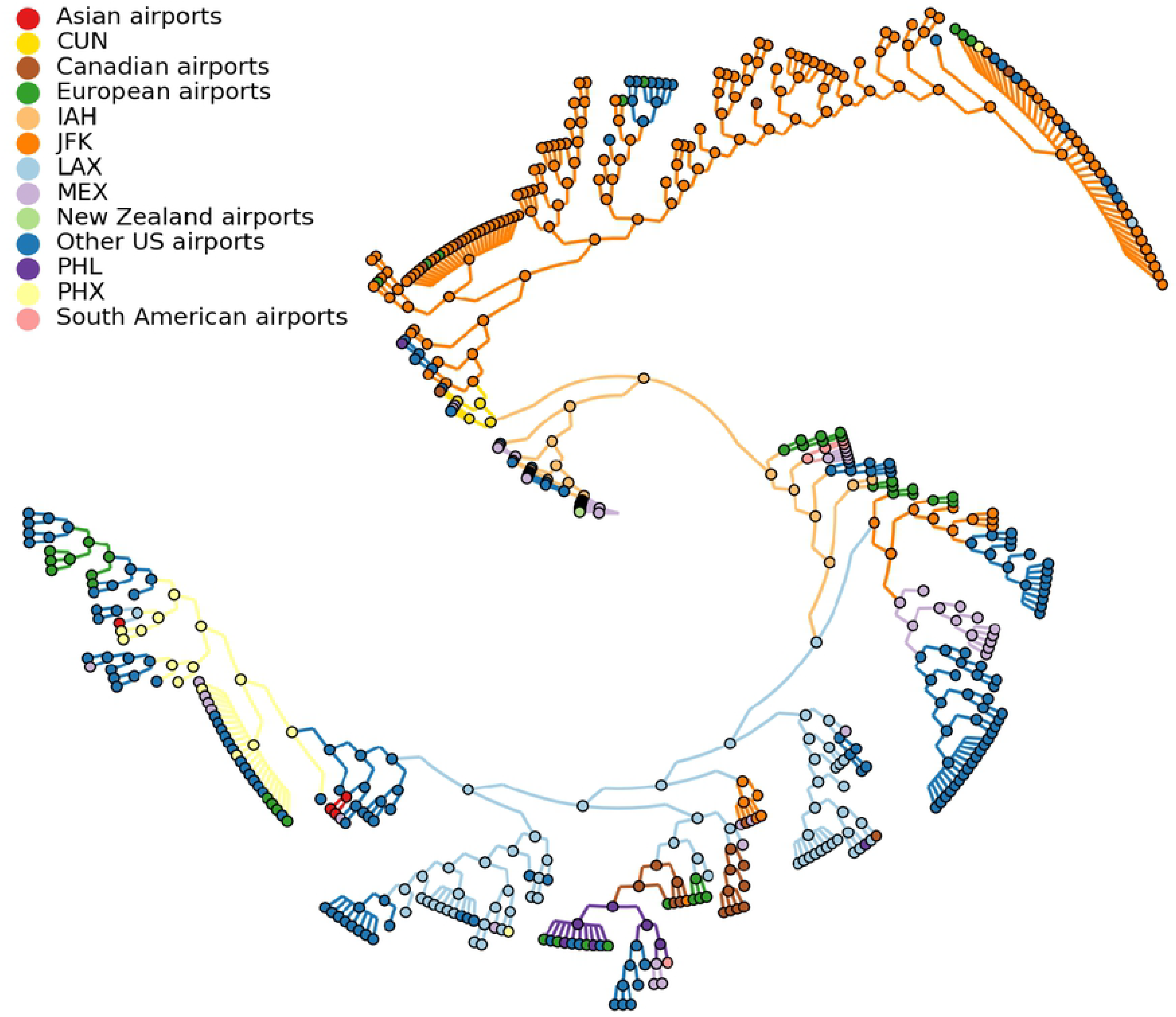
Phylogeographic reconstruction with the Sankoff algorithm using asymmetric, effective distances. Exemplary phylogeographic reconstruction using the Sankoff algorithm on the tree and the cost matrix shown in panel A. The cost matrix *c* is asymmetric and represents effective distances. For each internal node, the Sankoff algorithm calculates the minimal cost *S*(*i*) in the subtree, given the node is assigned location *i* (shown as the arrays in A, calculated via *S*(*i*) = *min*_*j*_[*c*_*ij*_ + *S*_*l*_(*j*)] + *min*_*k*_[*c*_*ik*_ + *S*_*r*_(*k*)], where *l* and *r* denote the two descendant subtrees, and their costs, *S*_*l*_(*j*) and *S*_*r*_(*k*), respectively. For the root (shown in red), location A results in the minimal cost and is assigned to that node (marked in green). Backtracking from the root to assign all other locations is shown in panel B. Given that a parent node has been assigned state *j*, the child node will be assigned the state *i* that minimizes *c*_*ji*_ + *S*(*i*). The result of the backtracking is indicated by arrows labeled with the costs and the states marked in green. The reconstructed spread along the tree is shown on a map in panel C.

An example of a reconstructed spread is shown next to the simulated spread in Fig 1B. In this reconstruction, the inferred origin was Zacatecas, a city north of Mexico City. From there, the virus spread to various locations including Mexico City and Cancún, which seeded a large number of infections in new locations. In the north of the USA, the virus further spread mainly via Chicago. In the simulated spread, locations like Mexico City and Chicago only play a minor role, but they could be unobserved intermediate locations due to connecting flights in the GLEAMviz simulation, where transitions are only reported via origin and destination. However, while the exact reconstructed paths differed from the simulated spread, most of the inferred internal locations in Mexico were geographically close. To quantify these geographic differences between spread paths, reconstructed phylogeographies were compared to the known spread on the simulated tree by calculating discrete Fréchet tree distances [13] using geographic distances between locations (Fig 3A). This method compares the paths of locations from the root to each leaf node, calculates discrete Fréchet distances [22] between them and corrects the distance for each node by the number of paths.

**Fig 3.**
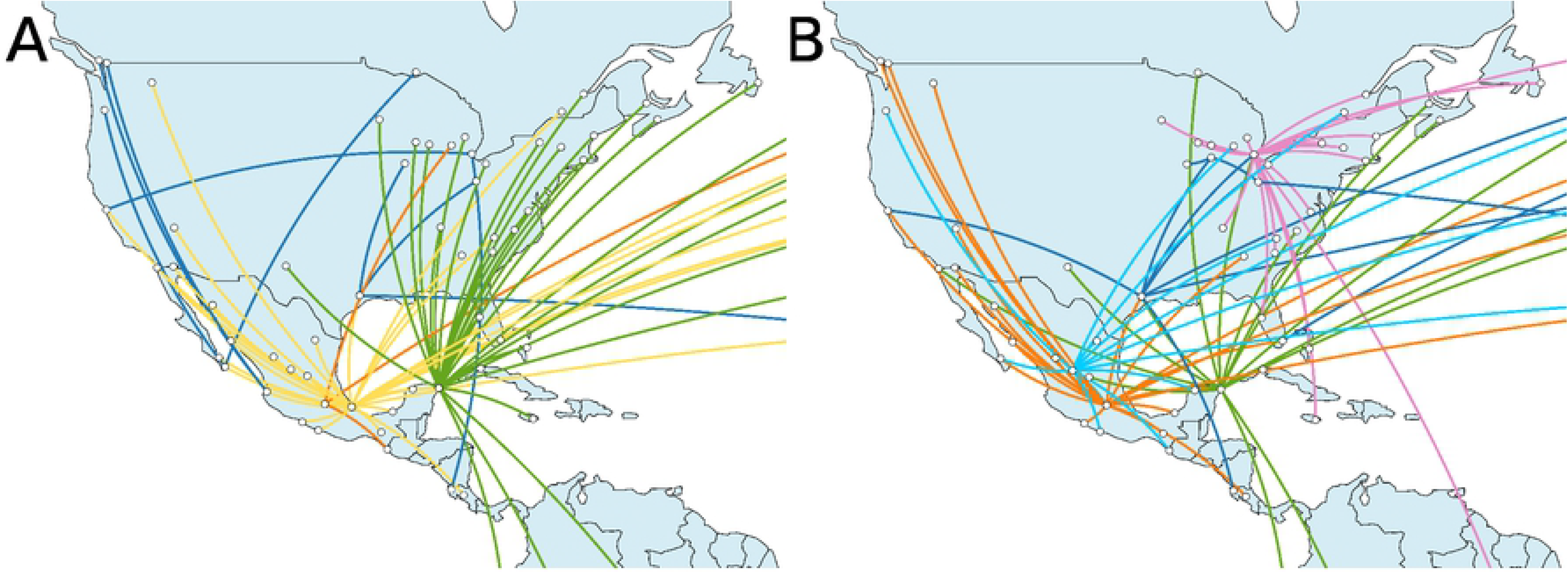
Parsimonious reconstructions with effective distances infer the simulated spread more accurately than reconstructions with geographic distances using BEAST. Boxplots of discrete Fréchet tree distances comparing simulated phylogeographies to reconstructed phylogeographies, using a total of 50 simulations. In panel A, the reconstructions use airports as locations and were performed on the tree inferred on simulated sequences using geographic (red) and effective (green) distances, and on the simulated tree, again using both geographic (blue) and effective (purple) distances. In panel B, the reconstructions use countries, the same way as described above. Additionally, reconstructions were inferred with BEAST with symmetric rates (green) and asymmetric rates (yellow).

For both the inferred and the simulated trees, the reconstruction using effective distances resulted in a lower distance compared to the reconstruction using geographic distances (P-values of paired t-test: 7.6 × 10^−10^ for inferred trees, 1.4 × 10^−10^ for reconstructed trees). However, both reconstructions with effective distances showed a small number of outliers with distances as high as in the analysis with geographic distances. When using geographic distances, the phylogeographic reconstruction using the inferred tree resulted in higher distances compared to the analysis on the simulated tree (P-value of paired t-test: 3.7 × 10^−9^), indicating that errors introduced by the tree inference influenced the results. When using effective distances, no significant difference was observed between reconstructions using the inferred and simulated tree (P-value of paired t-test: 0.1197). However, using the simulated tree resulted into a lower variance.

While the Fréchet tree distance measures distances between the entire spread routes, we further had a closer look at the inferred root locations. Since root locations indicate the possible origin of an outbreak, these are of particular interest. Veracruz, the correct root location in our simulation, wasn’t inferred except in two cases - once using geographic distances on the inferred tree, and once using effective distances on the simulated tree. However, the correct country of origin was inferred in the majority of cases when using effective distances. Interestingly, although no significant differences were observed when comparing the Fréchet tree distances, the reconstruction with the inferred tree topology inferred the country of origin less accurately (in 72% of the cases, compared to 98% on the simulated tree topology). When geographic distances were used, Mexico was only inferred as the origin in 24% of the simulations when using inferred topologies, and 34% when using simulated ones. Instead, the origin was placed in the US for most of the cases (74% and 62%, respectively).

Using the same 50 simulations as before, we repeated the analysis using countries instead of airports as locations. With this resolution, a comparison to the Bayesian reconstruction using BEAST was possible. As before, discrete Fréchet tree distances were calculated to compare the reconstruction to the reference data (Fig 3B). The parsimonious reconstruction using countries was comparable to the reconstruction using airports: using effective distances resulted in lower Fréchet tree distances than geographic ones (P-values of paired t-test: 1.5 × 10^−9^ for inferred trees, 5.6 × 10^−6^ for simulated trees). Fréchet tree distances were lower on reconstructions using the simulated tree topology as with the inferred one, but only when using geographic distances (P-values of paired t-test: 0.0231 for geographic distances, 0.3483 for effective ones). Phylogeographic reconstructions using BEAST with symmetric rates showed higher distances than the parsimonious reconstruction with effective distances, but were comparable to the reconstruction with geographic distances (P-values of paired t-test: 0.2668 for the comparison to geographic distances, 3.7 × 10^−5^ for the comparison to effective distances). Using asymmetric rates for the Bayesian reconstruction resulted in similar, but slightly lower distances.

For nearly all datasets and analyses, the root was placed either in Mexico or the US, with effective distances inferring Mexico more often (70% of the cases using the inferred tree topology and 96% on the simulated one) than geographic distances (34% and 62%, respectively). Bayesian analyses inferred Mexico as the origin in 46% of the datasets when using symmetric rates, and in 66% when using asymmetric rates.

### Phylogeographic reconstruction of the early spread of the pandemic H1N1 influenza A virus

To test the parsimonious reconstruction with effective distances on a real dataset, we analyzed the 378 HA sequences sampled from the beginning of the pH1N1 outbreak until the end of April 2009. For the phylogeographic reconstruction, locations were assigned to each sequence based on the sampling locations, as stated in the isolate name. The resolution of these locations varied between cities, states and countries. We assigned the main airport (as defined via the highest number of passengers) of the respective city, state or country to the sequence. Three cities did not have an airport and instead we used the geographically closest airport available in our list of locations. In total, this resulted in a set of 67 unique locations. The phylogeographic reconstruction using the Sankoff algorithm was performed using effective distances and is displayed in Fig 4.

**Fig 4.**
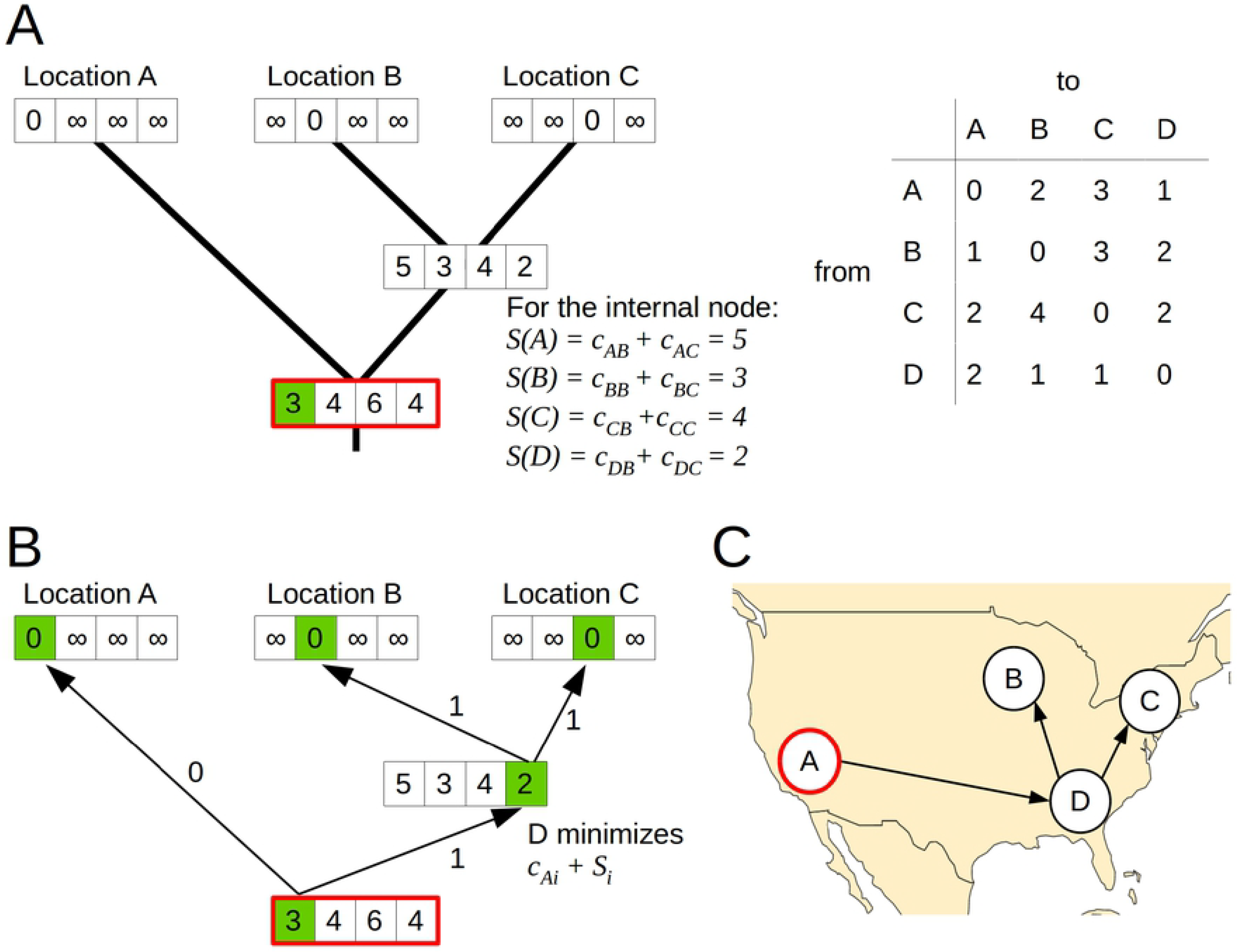
Phylogeny for pH1N1 viruses with reconstructed spread. Parsimonious phylogeographic reconstruction using effective distances on 378 HA sequences from the early stage of the 2009 H1N1 influenza A pandemic. Sequences from 67 different locations were included and represented by the closest airports. The main airports involved in the spread (IAH (Houston), LAX (Los Angeles), JFK (New York), PHL (Philadelphia), PHX (Phoenix)) are shown in separate colors, together with Cancun (CUN), which was not observed in the data but was inferred for some internal nodes. All other locations were summarized per country or continent to allow their visualization. The tree was visualized using GraPhlAn [23].

Our method inferred Mexico City for the root node of the tree and therefore as the origin of the outbreak. With that, we successfully reconstructed the correct country of origin. Mexico City is the main airport of the country and was assigned to all sequences sampled in Mexico due to the lack of more accurate geographic information. The actual suspected origin in La Gloria in the state Veracruz is around 300km away from our inferred origin. The lack of more precise information about the sampling locations likely prevents our method to infer the origin more closely. However, for some internal nodes our method inferred Cancún, a second location in Mexico. This demonstrates that our approach is able to infer intermediate locations which have not been observed at the tips and might therefore overcome some of the problems introduced by poor sampling resolution. Cancún is the second largest airport in Mexico and a common tourist destination. Therefore, it likely contributed to the global spread of the pH1N1 influenza A virus. People travelling from Cancún around mid April have been the first reported cases in the UK [24] and are further suspected to be the source of an outbreak at a school in New York [25]. Our reconstruction finds a link from Cancún to New York as well, together with links to Wisconsin and Alberta, Canada.

To explore the reconstructed spread in more detail, we counted the number of transitions to a new location for each observed location. 46 of the 68 locations only occured at terminal branches without seeding infections to new locations. Instead, there was a small number of airports mainly involved in the spread. The airports with the highest numbers of links to new locations were New York (31), Phoenix (24), Los Angeles (21), Houston (13), Mexico City (11) and Philadelphia (11). As expected with effective distances, the method infers large airports, but not only the largest airports of the region (as measured by the number of passengers) or the airports with the highest numbers of sequences (S1 Table). For example, Phoenix is the 10th largest airport in North America with 7 sampled sequences, Houston the 11th largest with 1 sequence, and Philadelphia the 20th largest with 4 sequences. This indicates that the main airports are not only determined by the numbers of passengers or the number of samples, but are dependent on the specific outbreak and its origin.

## Discussion

Parsimonious ancestral character state reconstruction has been used for phylogeographic inference in the past [6]. Instead of using the Fitch algorithm for the reconstruction, therefore minimizing the number of state transitions, we here used the Sankoff algorithm to minimize distances between locations along the tree. This allowed us to introduce two innovations: the inference of unobserved locations for internal nodes, as well as the direct inclusion of air transportation data via effective distances. While effective distances based on air travel are sensible for influenza A viruses, geographic distances are likely the best choice for pathogens with a local spread, including the historical spread of diseases or the spread of animal viruses like rabies. Effective distances can further be defined using local movements based on commuting data or gravity and radiation models [26,27], which can be useful to study epidemics in single countries where air travel does not play a major role. As long as prior knowledge about the mode of transportation is known, it’s simple to adjust the distance matrix used in the phylogeographic reconstruction with the Sankoff algorithm to a specific pathogen. To evaluate the phylogeographic reconstruction, we first simulated geographic spread using GLEAMviz and then the sampling, trees and sequences using FAVITES. It should be noted that the list of transmissions generated by GLEAMviz didn’t include transitions to locations that have been previously infected; an assumption which might be adequate for the very beginning of a pandemic but not in later stages. Further, the sampling process was simplified, with locations randomly chosen for sampling and only a small number of sequences per sampled location, while real influenza A virus sequences usually show a distinct geographic bias. Since the sequence diversity and tree resolution were similar to real pH1N1 sequence data and comparable locations were observed, we believe the simulation was appropriate for an evaluation of phylogeographic methods despite these simplifications. To our knowledge, no other reference dataset with known phylogeographic spread exists as an alternative.

By comparing the reconstructed spread paths to the simulated ground truth using discrete Fréchet tree distances, we showed that our method using effective distances inferred the phylogeographic spread more accurately than with geographic distances or the analysis with BEAST. However, when only considering the root instead of complete paths, the BEAST analysis with asymmetric rates inferred the correct country of origin more accurately than the reconstruction using geographic distances, and was comparable to the reconstruction with effective distances. The difference between effective and geographic distances indicated that it’s essential to choose suitable distances for the analysis. With suitable distances, accurate reconstructions could be achieved that outperformed the state-of-the-art. The parsimonious reconstruction further allowed to study viral spread in more detail than before. Instead of summarizing locations into large geographic areas like continents, the phylogeographic reconstruction was possible on fine-grained locations to the resolution of single cities. Both the simulated and real data proved that the analysis was feasible with a large set of total locations (3865 airports in the air transportation network) as well as observed locations; 76 locations on average - including 23 countries - in the simulated data and 67 in the real data. The reconstruction could be done within minutes, while the Bayesian reconstruction was time-consuming and ran for several days to reach a sufficient number of steps in the MCMC, even when using countries instead of airports. The parsimonious reconstruction using the Sankoff algorithm could further infer intermediate states not observed in the sample; a feature that was only achieved by continuous models so far.

However, the advantages of Bayesian methods to previous parsimonious methods still hold. Bayesian methods allow to integrate over uncertainties in the phylogeny and migration process, while parsimony methods assume a fixed tree topology that was previously inferred from the data and usually differs from the true tree topology. Parsimony methods further don’t infer posterior probabilities, therefore giving no indication about the certainty of the results, and do not consider branch lengths. When using the Sankoff algorithm, we assume prior knowledge about the mode of transportation, which might not always be available. Instead, Bayesian methods include the possibility to infer potential modes of transportation by testing predictors of spatial spread. In the future, both methods could be used in a complementary way depending on the data, the desired analysis as well as time and computing resources. To enable others to apply this approach to new datasets, all software is provided in a Github repository [28] and distance matrices are available on Zenodo [29].

The application to a dataset of HA sequences of the pH1N1 virus demonstrates how the parsimonious reconstruction using effective distances can be used in case of new outbreaks, as long as viral sequences and precise geographic information for the isolates are available. This approach offers a powerful tool to rapidly analyze sequence data, find the place of origin and propose possible spread routes to the resolution of single airports. This information would be helpful to implement control measures like increased surveillance or restricted travel to contain or slow down the global spread of a new infection.

## Methods

### Simulation

To create a reference dataset, we simulated the beginning of the pH1N1 influenza pandemic in 2009. We first simulated the geographic spread of the virus and then used these transition patterns to simulate isolate sampling, the phylogenetic tree and nucleotide sequences.

The geographic spread simulation was performed with GLEAMviz version 6.6 [15] using a stochastic SEIR (Susceptible-Exposed-Infectious-Recovered) model. We considered three compartments for infectious people: symptomatic (with travel), symptomatic (without travel) and asymptomatic, in line with previous studies [16,17]. In this model, the world is divided into 3252 metapopulations interconnected via an airport and commuting network. We set the origin of the pandemic to Veracruz, Mexico, on February 18th 2009, which was reported as the source and time of the outbreak [30] and has been used in similar models [17]. The simulation produced proportions of individuals in each SEIR compartment per day for each of the metapopulations, as well as the seeding location and day of first arrival of the disease for each newly infected population.

We used the latter as a list of transmissions to simulate the isolate sampling, the tree and the sequences. For these steps, the simulation software FAVITES was used [19]. To simulate the beginning of the pandemic, only the first 200 transition events were included, corresponding to day 85 in the simulation. We chose the number of samples per location via a Poisson distribution with *λ* = 0.5, therefore simulating an incomplete sampling of locations. The sampling times were chosen from an uniform distribution. Each sample represented one viral sequence in the subsequent analysis. Based on the sampling events, a tree was simulated using a coalescent model with exponential growth. Branch lengths on this tree corresponded to time as measured in days. We scaled these branch lengths with a rate of 0.000014, therefore assuming a rate of evolution of 0.00511 mutations per site per year, which is close to estimates reported for pH1N1 [31,32]. Given the tree with scaled branch lengths, nucleotide sequences of length 1700 were simulated under a GTR model using seq-gen [33]. This resulted in a set of sequences for each tip in the tree. These sequences along with the their sampling locations were then used for phylogeographic inference, while the simulated tree (including the locations for the internal nodes as given by the transition events) was used as a reference dataset.

### Data download

Nucleotide sequences of the hemagglutinin (HA) protein of pH1N1 were downloaded from the GISAID database [34]. All complete sequences from the beginning of the pandemic in March until end of April were downloaded, resulting in a dataset of 378 sequences.

### Phylogenetic reconstruction

For both simulated and real data, nucleotide sequences were aligned using MUSCLE version 3.8.31 with standard parameters [35]. Positions with gaps in more than 80% of the sequences were removed with TrimAl version 1.2 [36] to ensure a good quality of the alignment. Phylogenetic trees were then reconstructed with FastTree version 2.1.7 [37] using the GTR model. Simulated trees were rooted based on the ancestral sequence used in the simulation process, which was subsequently removed from the dataset. The tree based on real data was rooted using the sequence with the earliest sampling date.

### Parsimonious phylogeographic reconstruction

The phylogeographic reconstruction was based on the air transportation network from the OAG database including 3865 airports, which we used as all possible states in the analysis. Geographic distances between airports were calculated based on longitude and latitude using the R package geosphere [38]. This defined the distances as the shortest paths between locations, taking into account the ellipsoidal surface of the Earth. Effective distances were calculated based on the numbers of passengers travelling between airports in the year 2013, as in [11]. The air transportation network included airports from 228 countries, which were used as locations for the analysis on a country level. Geographic distances between countries were calculated as described above using the coordinates of the centroids. For effective distances, we aggregated the passenger numbers per country and recalculated the distances. The distance matrices were used as a cost matrix for the parsimonious reconstruction using the Sankoff algorithm [12]. Distances represent the cost of traveling from one state to another and the Sankoff algorithm finds the internal states with the minimal cost, i.e. minimizing the distance the virus travelled along the tree. An example of the inference using asymmetric distances is shown in Fig 1. We used delayed transformation in case of ambiguities. When an ambiguity occured at the root and therefore couldn’t be resolved with delayed transformation, we randomly chose one of the possible states. Since effective distances are generally not symmetric, the tree was rooted before the reconstruction.

### Bayesian phylogeographic reconstruction

The simulated data included a large number of states with relatively few sequences, which is not suitable for Bayesian phylogeographic reconstruction [9]. Using countries instead of airports reduced the number of states to make the Bayesian reconstruction feasible. We used BEAST version 2.4.8 for this analysis [14]. Sampling dates were included into the phylogenetic reconstruction as measured in days. A HKY nucleotide substitution model was used together with a strict molecular clock. For the tree prior, a coalescent model with exponential population growth was chosen. For the phylogeographic reconstruction, analyses were performed separately with both symmetric and asymmetric rates. Markov Chain Monte Carlo (MCMC) was run for 100 million steps with trees sampled every 10,000 steps, resulting in a sample of 10,001 trees.

Tracer version 1.7.1 [39] was used to confirm adequate effective sample sizes (ESS), indicating good estimates of the posterior distributions of the parameters. TreeAnnotator was then used to summarize the sampled trees into a maximum clade credibility tree using a burn-in of 10%. For the evaluation of the phylogeographic reconstruction, we assigned the location with the highest posterior probability to each node.

## Acknowledgements

We thank Dirk Brockmann for providing the air passenger data to calculate effective distances between airports.

## Supporting Information

**S1 Fig. Comparison of real and simulated data**.

A) Comparison of pairwise genetic distances between sequences for both real HA sequence data and the simulated sequences. B) Comparison of branch lengths on trees inferred on both real HA sequence data and simulated sequences.

**S1 Table. Main airports involved in the spread of pH1N1**.

Main airports involved in the spread of pH1N1, as measured by the number of transitions to new locations. The airports are New York (JFK), Phoenix (PHX), Los Angeles (LAX), Houston (IAH), Mexico City (MEX) and Philadelphia (PHL). The size of the airports in North America was determined by the numbers of passengers in 2013 as given via the OAG database.

